# Countries are out of step with international recommendations for tuberculosis testing, treatment, and care: Findings from a 29-country survey of policy adoption and implementation

**DOI:** 10.1101/533851

**Authors:** Karishma Saran, Tiziana Masini, Isaac Chikwanha, Gregory Paton, Rosalind Scourse, Patricia Kahn, Suvanand Sahu, Christophe Perrin, Lucica Ditiu, Sharonann Lynch

## Abstract

**Background:** Tuberculosis (TB) poses a global health crisis requiring robust international and country-level action. Adopting and implementing TB policies from the World Health Organization (WHO) is essential to meeting global targets for reducing TB burden. However, many high TB burden countries lag in implementing WHO recommendations. Assessing the progress of implementation at national level can identify key gaps that must be addressed to expand and improve TB care.

**Methods:** In 2016/2017, Médecins Sans Frontières and the Stop TB Partnership conducted a survey on adoption and implementation of 47 WHO TB policies in the national TB programs of 29 countries. Here we analyze a subset of 23 key policies in diagnosis, models of care, treatment, prevention, and drug regulation to provide a snapshot of national TB policy adoption and implementation. We examine progress since an analogous 2015 survey of 23 of the same countries.

**Results:** At the time of the survey, many countries had not yet aligned their national guidelines with all WHO recommendations, although some progress was seen since 2015. For diagnosis, about half of surveyed countries had adopted the WHO-recommended initial rapid test (Xpert MTB/RIF). A majority of countries had adopted decentralized models of care, although one-third of them still required hospitalization for drug-resistant (DR-)TB. Recommended use of the newer drugs bedaquiline (registered in only 6 high-burden TB countries) and delamanid (not registered in any high-burden country) was adopted by 23 and 18 countries, respectively, but short-course (9-month) and newer pediatric regimens by only 13 and 14 countries, respectively. Guidelines in all countries included preventive treatment of latent TB infection for child TB contacts and people living with HIV/AIDS, but only four extended this to adult contacts.

**Conclusion:** To reach global TB targets, greater political will is needed to rapidly adopt and implement internationally recognized care guidelines.

**KEY MESSAGES:** *What is already known?:*

- Countries may be slow to adopt and implement updated World Health Organization (WHO) Tuberculosis (TB) testing, treatment, and prevention recommendations.
- Implementing updated TB guidelines from WHO is a fundamental first step to honoring international commitments, made through the United Nations (UN) Sustainable Development Goals (SDGs) and UN High-Level Meeting on TB Political Declaration, to end TB by 2030.

*What are the new findings?:*

- Of 29 mostly high TB burden countries, none had fully aligned their national guidelines with WHO recommendations, although some progress has been made since 2015.
- A lack of alignment with WHO recommendations was found across all policy areas surveyed, including prevention, diagnosis, treatment, models of care and drug regulation, particularly regarding uptake of newer, faster, more effective approaches.

*What do the new findings imply?:*

- To reach global TB targets, greater political will is needed to adopt and implement internationally recognized care guidelines more rapidly, and specifically, to keep up with the latest recommendations.
- Periodic surveys of progress at the national level are a valuable way to identify specific areas where countries or regions have fallen behind and that require specific policy and/or programmatic attention.

## INTRODUCTION

Over the past decade, progress against the global tuberculosis (TB) crisis has slowed, with TB becoming the world’s leading single infectious disease killer, accounting for 1.6 million deaths in 2017.^1^ Although TB is preventable and treatable, global efforts to end the epidemic have been off target according to the World Health Organization (WHO)’s annual TB reports over the last 7 years – an estimated 42 million people out of nearly 67 million were diagnosed and put on treatment, missing close to 25 million people with TB.^1, 2, 3, 4, 5, 6, 7^ In 2017, an estimated 10 million people were infected with TB, but almost 40% of them, and 75% of the 558,000 people with drug-resistant (DR)-TB, were missed from diagnosis and treatment.^1^

A handful of newer, more effective health tools and improved strategies of care have been developed in the past several years and recommended by WHO. These newer tools and strategies range in modality from diagnosis and treatment, to prevention and patient-centered models of care. Through the United Nations (UN) Sustainable Development Goals (SDGs)^8^ and UN High-Level Meeting on TB Political Declaration,^9, 10^ countries have committed to ending TB by 2030, and a critical first step is to ensure national TB guidelines reflect WHO policy recommendations. However, countries with considerable TB burdens have been slow to adopt and implement WHO recommendations on the use of newer tools and strategies, hindering access for people affected by TB who need them.

For the first time in nearly half a century, new tools that substantially improve TB diagnosis and treatment have become available in the past several years. Newer diagnostic tests, such as Xpert MTB/RIF, and two newer drugs for treating DR-TB, bedaquiline (BDQ) and delamanid (DLM), have been recommended by WHO in the last decade.^11^ These innovations are desperately needed; using the decades-old standard treatment, the cure rate for multidrug-resistant (MDR)-TB is 55%, dropping to 34% for those with extensively drug-resistant (XDR-)TB.^1^ While evidence accumulates showing that BDQ and DLM improve treatment outcomes, these newer drugs remained inaccessible to nearly 90% of people eligible to receive them in 2017, based on WHO recommendations at the time.^12^

Improved strategies in TB prevention and models of care are also recommended internationally. For prevention, WHO recommends testing and treatment of latent TB infection in high-risk populations,^13^ as well as other strategies such as active case finding. The first pillar of the WHO End TB Strategy is “Integrated, patient-centered TB care and prevention”, in which decentralized, community-based models of care are recommended as the preferred treatment strategy.^14^ Another important model of care is integrated treatment of TB and HIV for people living with or at risk for either disease.

From 2014 to 2017, Médecins Sans Frontières (MSF) and the Stop TB Partnership (STBP) conducted three surveys on the status of key TB policies and related practices in high-burden countries, to monitor progress and identify gaps in the adoption of WHO recommendations into national TB programs.^15, 16, 17^ Called the “Out of Step” (OOS) surveys, these reports were comprehensive, with the latest report examining 47 policy indicators in five areas – diagnostics, models of care, treatment, prevention, and drug regulation – across 29 countries.^17^

Here we present the findings of a subset of key policy and related practice indicators from the most recent survey and discuss their implications on access to diagnosis and treatment for people with TB. As our survey was conducted before new 2018 WHO DR-TB treatment recommendation updates, we provide here a snapshot of the state of national TB policy adoption and levels of care compared to WHO recommendations at the time of study in late 2016 to early 2017.

## METHODS

A detailed description of the methodology can be found in the Supporting Information.

### Country Inclusion

The 2017 OOS survey was conducted in 29 countries between October 2016 and May 2017, which together were home to 82% of the global TB burden (Table 1). These countries were selected based on having a high burden of TB, MDR-TB, and/or TB/HIV co-infection according to WHO criteria^18^ and/or an MSF or STBP TB project presence. For 2016-2020, each of three WHO high-burden country lists (TB, MDR-TB, and TB/HIV) is defined as the top countries worldwide in absolute numbers of cases, plus an additional 10 countries not already on the list and that have the highest case rates per capita; and that meet a minimum threshold in terms of absolute numbers of cases (10,000 per year for TB, and 1,000 per year each for MDR-TB and TB/HIV).

**Table 1.** Countries Surveyed

| Country | Diagnosis Gap<br>% of people with estimated<br>TB (new and relapse) missing<br>in the country's case<br>detection rates | DR-TB Treatment Gap<br>% of people with DR-TB<br>not yet receiving<br>treatment, relative to<br>incidence |
| --- | --- | --- |
| Afghanistan* | 30 | 94 |
| Armenia* | 20 | 65 |
| Bangladesh | 33 | 89 |
| Belarus | 20 | 41 |
| Brazil | 13 | 59 |
| Cambodia | 34 | 88 |
| Central African Republic | 51 | 43 |
| China | 13 | 92 |
| Democratic Republic of the Congo | 43 | 89 |
| Eswatini (formerly Swaziland) | 20 | 48 |
| Ethiopia | 32 | 88 |
| Georgia* | 23 | 47 |
| India | 35 | 71 |
| Indonesia | 47 | 87 |
| Kazakhstan | 0 | 14 |
| Kenya | 47 | 86 |
| Kyrgyzstan | 23 | 67 |
| Mozambique | 48 | 89 |
| Myanmar | 32 | 81 |
| Nigeria | 76 | 93 |
| Pakistan | 32 | 89 |
| Papua New Guinea | 26 | 80 |
| Philippines | 45 | 79 |
| Russian Federation | 2 | 48 |
| South Africa | 32 | 23 |
| Tajikistan | 22 | 64 |
| Ukraine | 26 | 51 |
| Vietnam | 17 | 62 |
| Zimbabwe | 29 | 81 |

\*At the time of the survey, all countries were WHO-defined as high burden for TB, MDR-TB, or TB/HIV co-infection, except for Afghanistan, Armenia, and Georgia.
Source: WHO, 2017 (http://www.who.int/tb/country/data/profiles/en/)

### Policy Adoption Questionnaire

A semi-structured questionnaire was developed between September and November 2016, to assess the national adoption and implementation of 47 WHO TB policies and related practices in five key areas: diagnostics; models of care; treatment regimens for drug-sensitive (DS) and DR-TB; prevention; and drug regulatory environment. The policies included in the survey were selected based on their importance reflecting a relatively recent optimization or improvement over a previous policy or practice; policies and practices where uneven adoption across countries was observed; or, in the case of national drug regulatory environment, practices that are essential for bringing newer medicines and recommended drug regimens into routine use.

The questionnaire asked whether national policies were aligned with (had adopted) current WHO policies, calling for yes or no answers; in cases of no response or unclear answers, the answer was recorded as “unknown”. If yes, then the questionnaire asked if these policies had been implemented or not. An additional box for each question asked the respondents to identify the specific policy document used to support their answer.

In this study, we analyzed a subset of 23 key policy and related practice indicators, of the 47 policies and practices included in the survey questionnaire, that impact access to TB care for people affected by TB.

### Data Collection and Validation

Between October and November 2016, MSF and STBP conducted a desktop review of relevant TB and HIV policy documents and guidelines from each country. Managers of NTPs were contacted and asked to review the documents to ensure they reflected the most up-to-date policy guidance at the time. NTPs were given approximately three weeks to confirm if these documents corresponded to the latest policy guidance being used, or to send any additional documents. This process continued until the end of December 2016. The full list of documents obtained is available in the online version of the OOS report.^17^

For the completion of the questionnaires, MSF followed up in the 18 countries where it operates TB projects, and STBP followed up in the remaining 11 countries. MSF teams in country were contacted and asked to complete the questionnaire (December 2016 to January 2017), while STBP pre-filled the questionnaire using the collected policy documents and asked the NTP in each country to validate the responses (February to May 2017).

Once the questionnaires were completed, each one was reviewed by MSF experts and a consultant fact-checker. Extensive efforts were made to follow up with respondents by e-mail or phone in order to verify or clarify responses where necessary, including if one or more of the documents mentioned in the responses was missing or we could not identify the source of the information.

### Data Analysis

The findings presented here are reported as both percentages and numbers. Unless noted, the denominator is 29, for all countries included in the survey. If a country did not answer a question, both the numerator and denominator were adjusted.

Also, progress made in the 2017 survey compared to the 2015 OOS survey are presented where possible for the 23 countries included in both surveys, reported as both percentages and numbers. Only data that could be verified with no discrepancies were included in this comparison, so for some parameters the denominators were less than 23.

## RESULTS

### Diagnosis

To ensure people with TB are treated appropriately, and to prevent further transmission, diagnosis needs to be quick and accurate. The WHO End TB Strategy calls for all countries to implement initial diagnostic testing with a WHO-recommended rapid diagnostic test by 2020.^19^

Starting in 2010, WHO recommended the Xpert *Mycobacterium tuberculosis* (MTB)/rifampicin (RIF) resistance test as the initial diagnostic test for all suspected cases of TB and MDR-TB, replacing microscopy.^20^ Our findings showed that Xpert MTB/RIF was adopted in policy as the initial test for all suspected TB cases by only about half (52%, 15/29) of the countries surveyed (Figure 1). This was an improvement on the 2015 OOS survey, when only 32% (7/22) of countries had adopted the use of Xpert MTB/RIF as an initial test in their policies, compared to 68% (15/22) of these same countries in 2017 (Figure 2).

**Fig 1.**
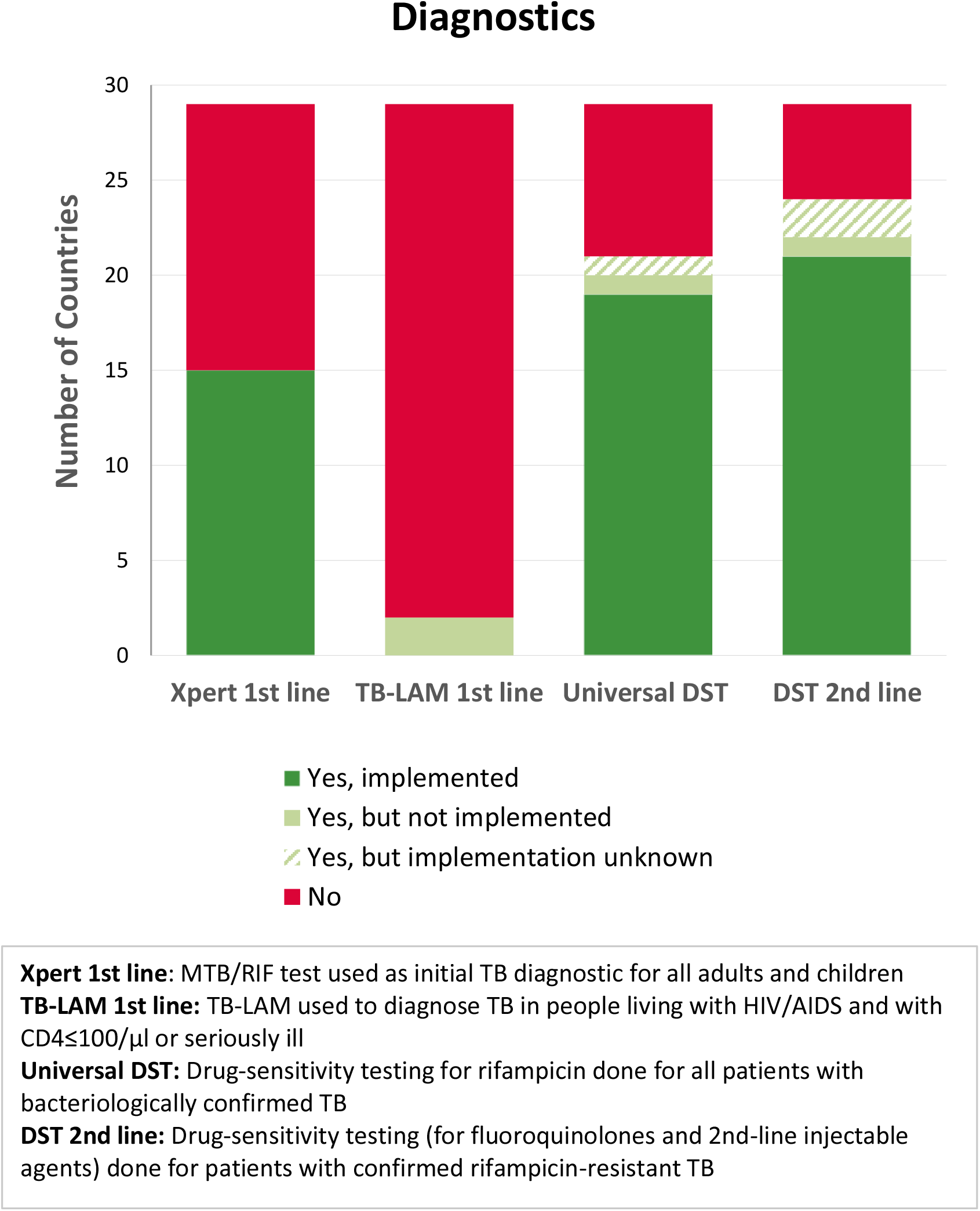
Diagnosis policy adoption and implementation

**Fig 2.**
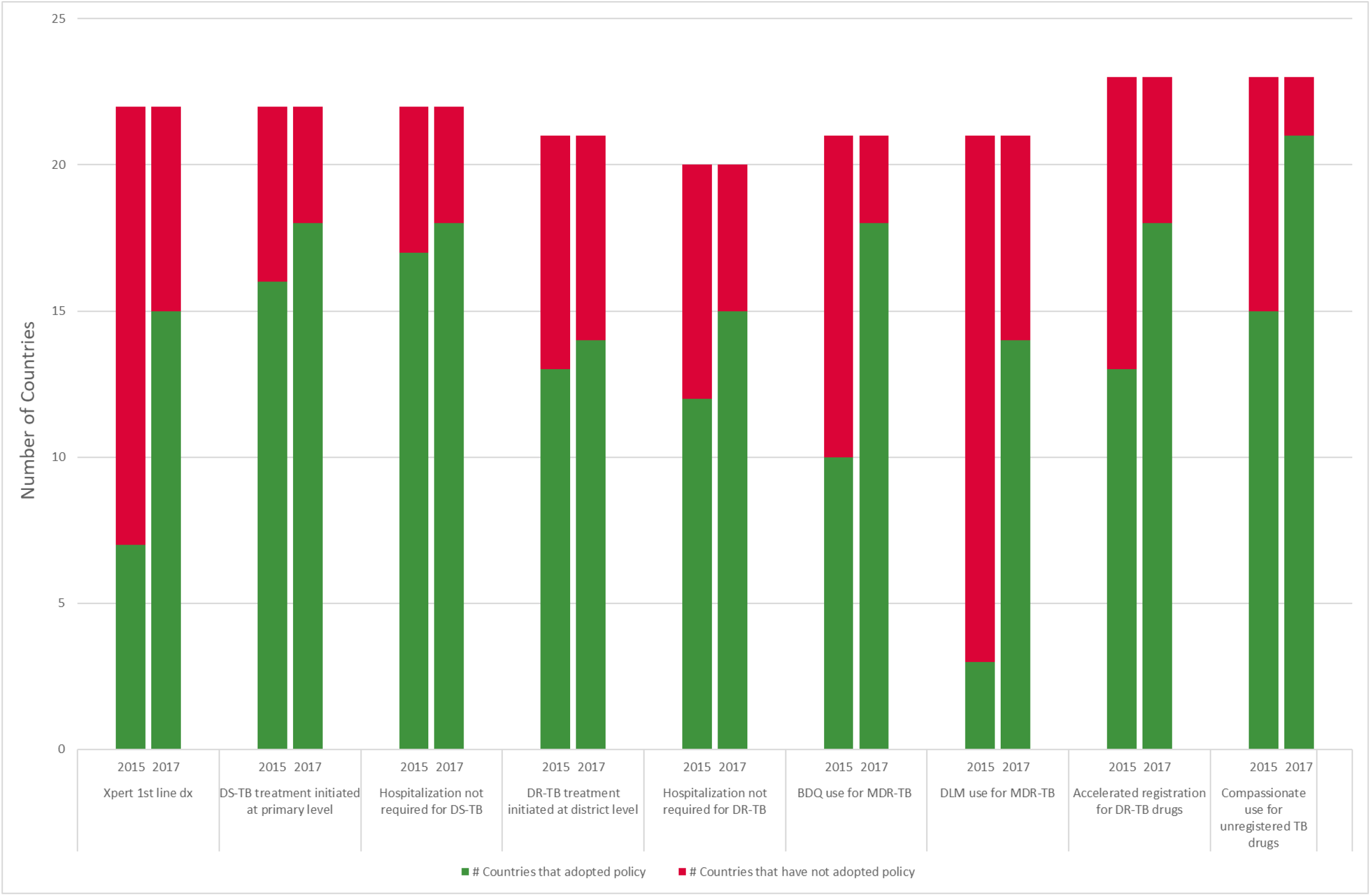
Changes in adoption of policies and related practices, 2015 to 2017

Due to specific challenges in diagnosing TB in people living with HIV/AIDS, in 2015 WHO recommended the use of TB lateral flow urine lipoarabinomannan assay (TB-LAM) to assist the diagnosis of TB for co-infected people with CD4 cell counts ≤100 cells/μl plus TB symptoms, or those who are very ill.^21^ We found that only two countries had adopted a guideline reflecting this recommendation, but neither had implemented it (Figure 1). Three other countries used TB-LAM in selected facilities, while two others used it for research purposes only.

The WHO End TB Strategy includes detecting all cases of DR-TB through drug-susceptibility testing (DST), calling on countries to adopt universal DST for at least rifampicin resistance (RIF-DST), an important first-line drug, for all people with bacteriologically confirmed TB; and subsequently, DST for at least fluoroquinolones (FLQ) and second-line injectable agents (SLIA), for all TB patients with confirmed rifampicin-resistant-TB (RR-TB). Albeit relatively new recommendations at the time of this survey, RIF-DST for all people with bacteriologically confirmed TB was in the guidelines of 72% (21/29) of countries, and implemented by 19 of these 21 countries. Second-line DST for FLQs and SLIAs for at least all people with RR-TB were in the guidelines of 83% (24/29) of countries, and implemented by 21 of these 24 countries (Figure 1).

### Models of Care

WHO recommendations in 2017 stated that community-or home-based treatment is the preferred option for people on TB treatment, and that decentralized models of care are recommended for DR-TB patients, unless they are extremely ill.^14^ The healthcare facility level at which TB treatment can be initiated, as well as the requirement of hospitalization for treatment, are two indicators that can be looked at to assess the extent to which countries have implemented decentralized models of care.

Decentralized initiation of DS-TB treatment was relatively widespread among the countries surveyed, with some progress since 2015 (Figures 2–3). The survey showed that DS-TB treatment can be started at the primary health care level in 83% (24/29) of countries, and has been implemented in 22 of these 24 countries. This policy adoption of decentralized DS-TB treatment increased from 73% (16/22) in 2015 to 82% (18/22) of the same countries in 2017. However, routine hospitalization for the initiation or some portion of treatment of DS-TB was still required in 21% (6/29) of countries.

**Fig 3.**
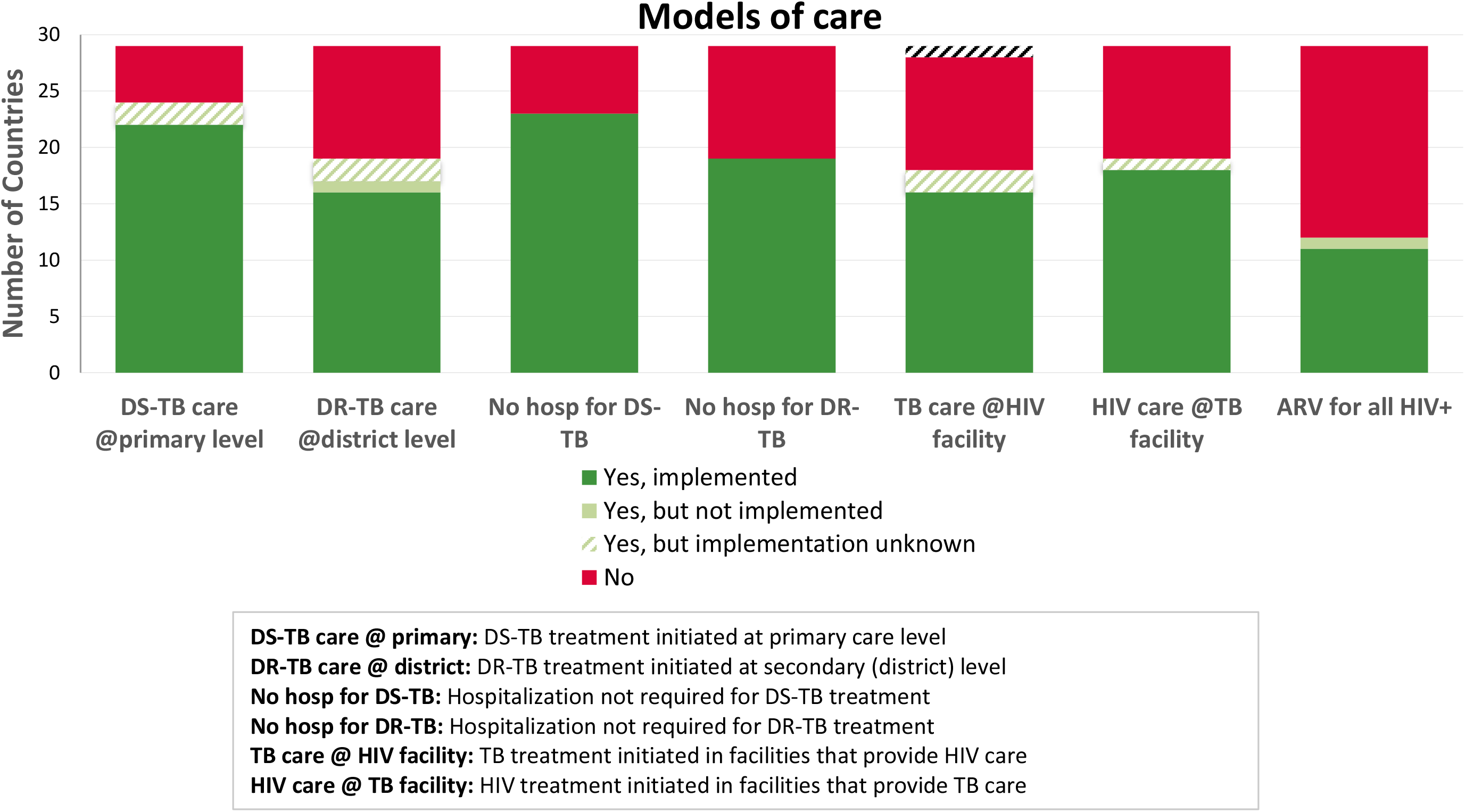
Models of care policy and related practice adoption and implementation

This had decreased slightly since 2015, when 23% (5/22) of countries required routine hospitalization, compared to 18% (4/22) of the same countries in 2017.

Countries were found to be slower to adopt decentralized models of care for patients with DR-TB, with lower adoption and implementation rates and minimal progress from 2015 to 2017 (Figures 2–3). DR-TB treatment at the district level was recommended in 66% (19/29) of the countries, 16 of which had implemented the policy at some level. In comparison, in 2015, DR-TB treatment could be initiated at the district level in 62% (13/21) of countries, increasing by only one country to 67% (14/21) of the same countries in 2017. Routine hospitalization for the initiation or some portion of treatment of DR-TB was also still required in 35% (10/29) of the countries, with minimal improvements since 2015 when 25% (5/20) of countries required routine hospitalization for DR-TB patients, compared to 40% (8/20) of the same countries in 2015.

Another key model of care is integrated TB/HIV treatment. TB is the leading cause of death among people living with HIV/AIDS (PLWHA), and PLWHA are 26-31 times more likely to develop active TB.^22^

Since 2015, WHO has recommended antiretroviral therapy (ART) for all PLWHA,^23^ called the “Test and Start” approach. Given the benefits of this approach in reducing morbidity and mortality due to TB, implementation of this policy is an urgent step for countries with a high burden of HIV/TB co-infection. Our survey found that WHO’s “Test and Start” policy had been adopted by only 41% (12/29) of countries, and 11 of these countries had implemented the policy (Figure 3). Furthermore, despite being recognized and recommended by the WHO since the previous six years, TB treatment provision in HIV care facilities was in the policies of only 64% (18/28) of countries, and HIV treatment provision in TB care was a policy in only 66% (19/29) of countries.

### Treatment

Bedaquiline (BDQ) and delamanid (DLM) are the two most recent TB drugs to be recommended by WHO, offering less toxic and more effective treatment courses. BDQ and DLM were first recommended for MDR-TB by WHO in 2013^24^ and 2014,^25^ respectively. In January 2018, WHO updated the guidance around the use of DLM,^26^ and in August 2018, updated guidance on the use of BDQ for MDR-TB treatment.^27^ Our survey was conducted prior to these newer guidelines (see Discussion).

At the time of the survey, BDQ was included in the national guidelines for DR-TB treatment in 79% (23/29) of countries, while DLM was included in the guidelines in 62% (18/29) of countries (Figure 4). This was an improvement from 2015, when BDQ was included in 48% (10/21) of countries compared with 86% (18/21) of the same countries in 2017 (Figure 2). DLM was included in national guidelines in only 14% (3/21) in 2015, but this rose significantly to 67% (14/21) for these countries in 2017 (Figure 2). However, in the 2017 survey, only 17/23 countries for BDQ and 11/18 for DLM stated the policy had actually been implemented at any level (Figure 4).

**Fig 4.**
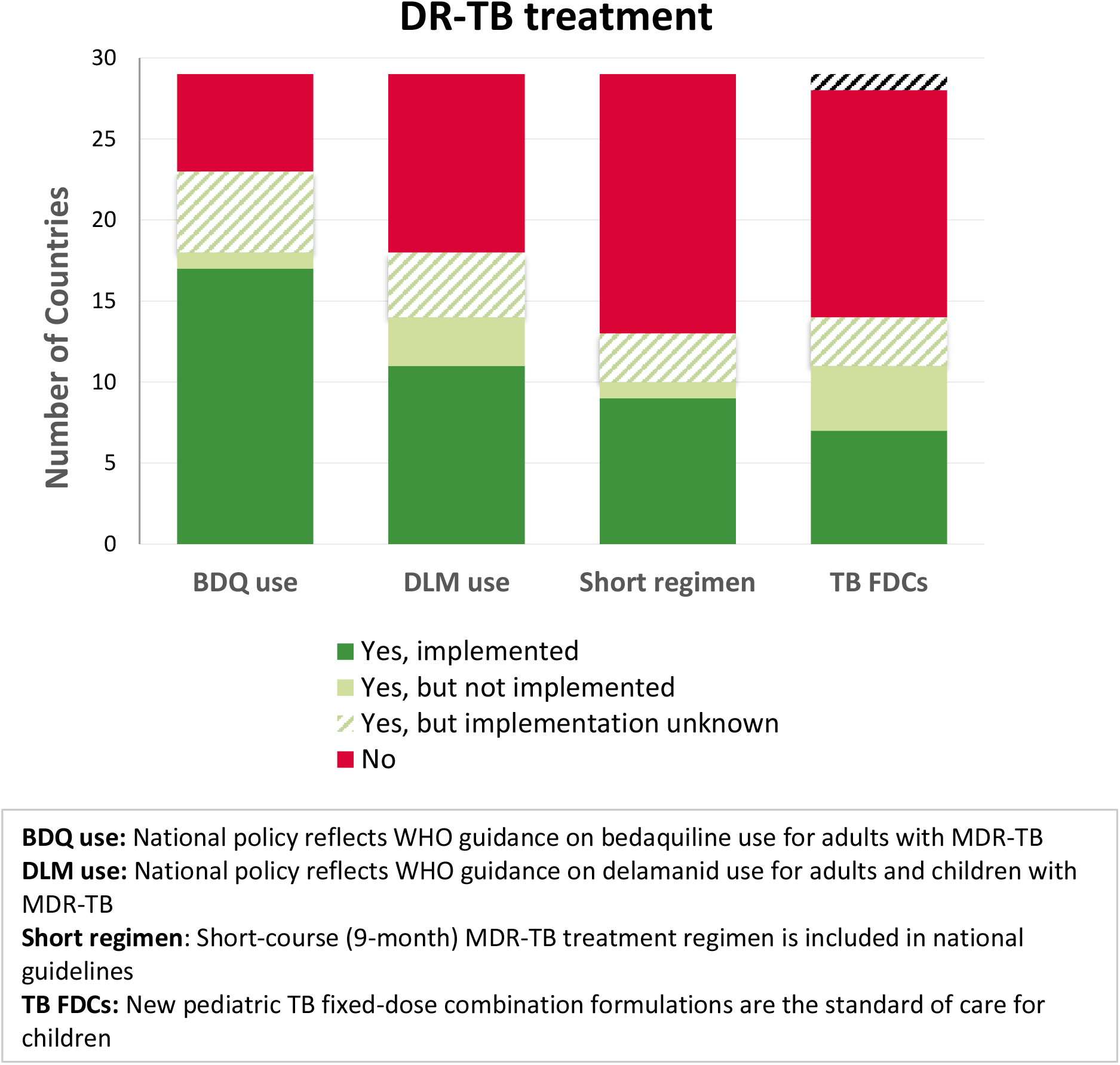
Treatment policy adoption and implementation

In 2016, the WHO-recommended a shorter treatment regimen for rifampicin resistant (RR)-or MDR-TB using existing drugs but reducing the treatment duration from 20 months to 9-12 months.^28^ By the time of the survey, adoption and implementation of this regimen had been minimal, but the recommendation was also relatively new at that time. We found that the 9-month MDR-TB treatment regimen was included in the guidelines in only 45% (13/29) of the countries surveyed; and only 9 countries had implemented this recommendation (Figure 4).

The lack of child-friendly drug formulations complicated pediatric TB treatment until December 2015, when new fixed-dose combinations (FDCs) of pleasant-tasting, WHO-recommended pediatric drug formulations for DS-TB became available.^29^ Despite transforming TB treatment for children, the new pediatric TB FDCs were the standard of care in only 50% (14/28) of countries, and were implemented in only 7 of those countries (Figure 4).

### Prevention

Prevention is essential to achieving the goals and targets of the End TB Strategy. The Strategy calls for the expansion of preventive treatment, also known as treatment for latent TB infection (LTBI), of people with a high risk of TB. At the time of the survey, WHO recommended systematic testing and treatment of LTBI for PLWHA, and adult and child contacts of pulmonary TB.^13^

We found favorable adoption of policy but inadequate implementation. Countries prioritized LTBI treatment in children <5 years old and PLWHA. According to national policy, all countries (29/29) surveyed provided LTBI treatment to child contacts <5 years old and PLWHA, with the policy implemented in 83% (24/29) of them. Fourteen percent (4/29) of countries had the policy for LTBI treatment for adult contacts (Figure 5). At the time of the survey, WHO had recommended LTBI treatment for children <5 years old and PLWHA, but not for adult contacts, which was recommended later in 2018; these 4 countries were ahead of the WHO recommendation.

**Fig 5.**
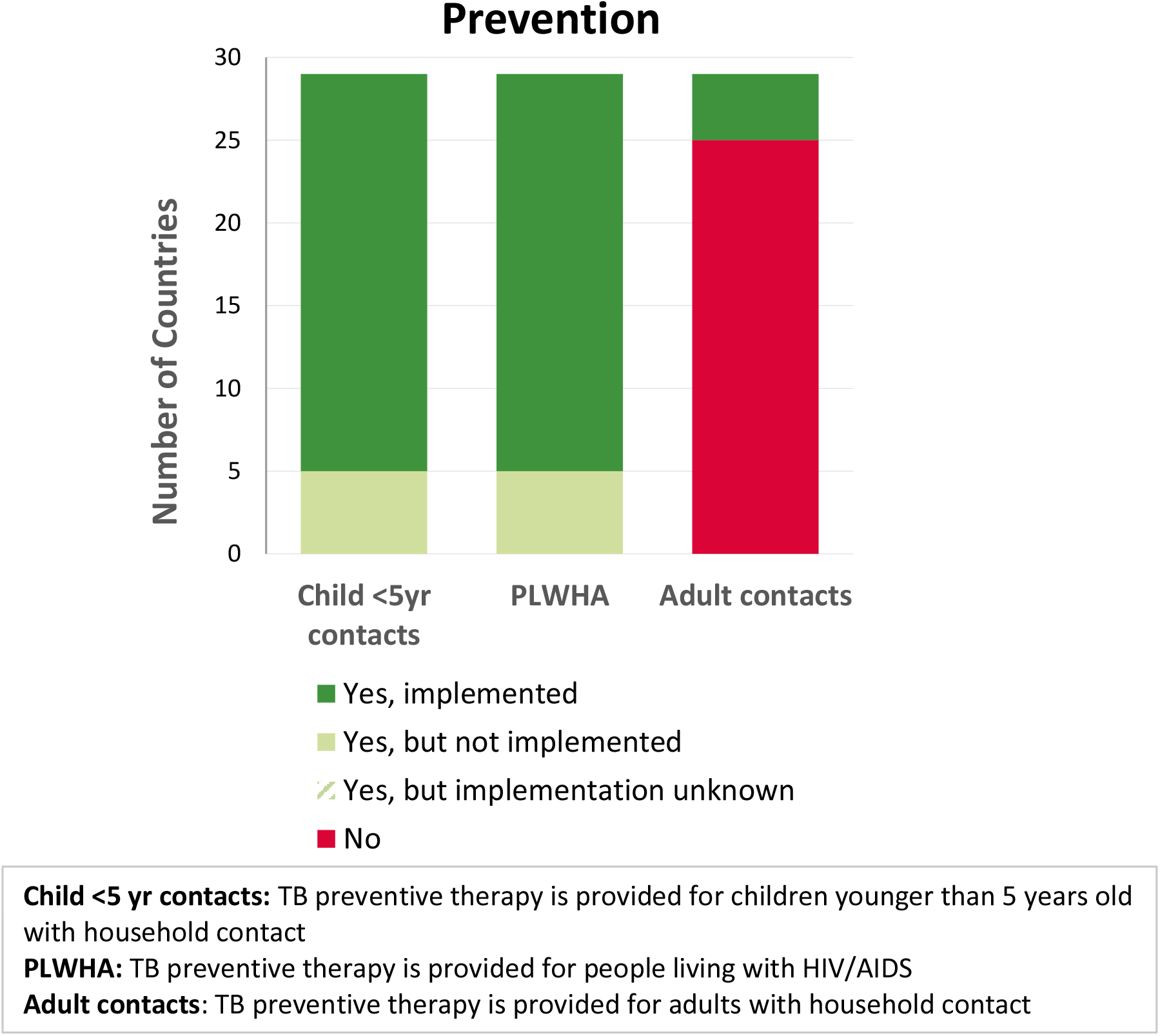
Preventive latent TB treatment policy adoption and implementation

### Drug Registration

Treatment for people affected by TB is dependent on access to medicines. New and repurposed TB drugs need to be registered in each country that needs them by the manufacturers. In countries where they have not yet been registered, alternative access mechanisms are needed to expedite registration of quality-assured sources, such as accelerated registration of WHO-prequalified and Stringent Regulatory Authority registered medicines and the WHO Collaborative Registration Procedure (CRP). Other mechanisms such as compassionate use programs or import waivers can also provide short-term access solutions to unregistered medicines. Although these mechanisms are not WHO policies per se, they are related practices indicative of national TB treatment access.

At the time of the survey, manufacturers had not registered BDQ or DLM in many TB high-burden countries. The survey showed BDQ was only registered in six of the countries, while DLM was not registered in any of the 29 countries (Figure 6). Seventy-five percent (21/28, one country’s status was unknown) of the countries reported having accelerated registration mechanisms in place that could be applied to new and repurposed DR-TB medicines. Among the 23 countries surveyed on this indicator in both the 2015 and 2017 surveys, 56% (13/23) of countries had mechanisms in place in 2015, increasing to 78% (18/23) in 2017 (Figure 2). Only 41% (12/29) of countries had adopted the practice of enrollment in the WHO CRP (Figure 6), and 6 countries had no accelerated registration mechanism in place and were not enrolled in the WHO CRP. Compassionate use or other national-level legal mechanisms were in place in 89% (25/28, one country’s status was unknown) of countries surveyed. In 2015, this was the case for 65% (15/23) of countries, increasing to 91% (21/23) in 2017 (Figure 2).

**Fig 6.**
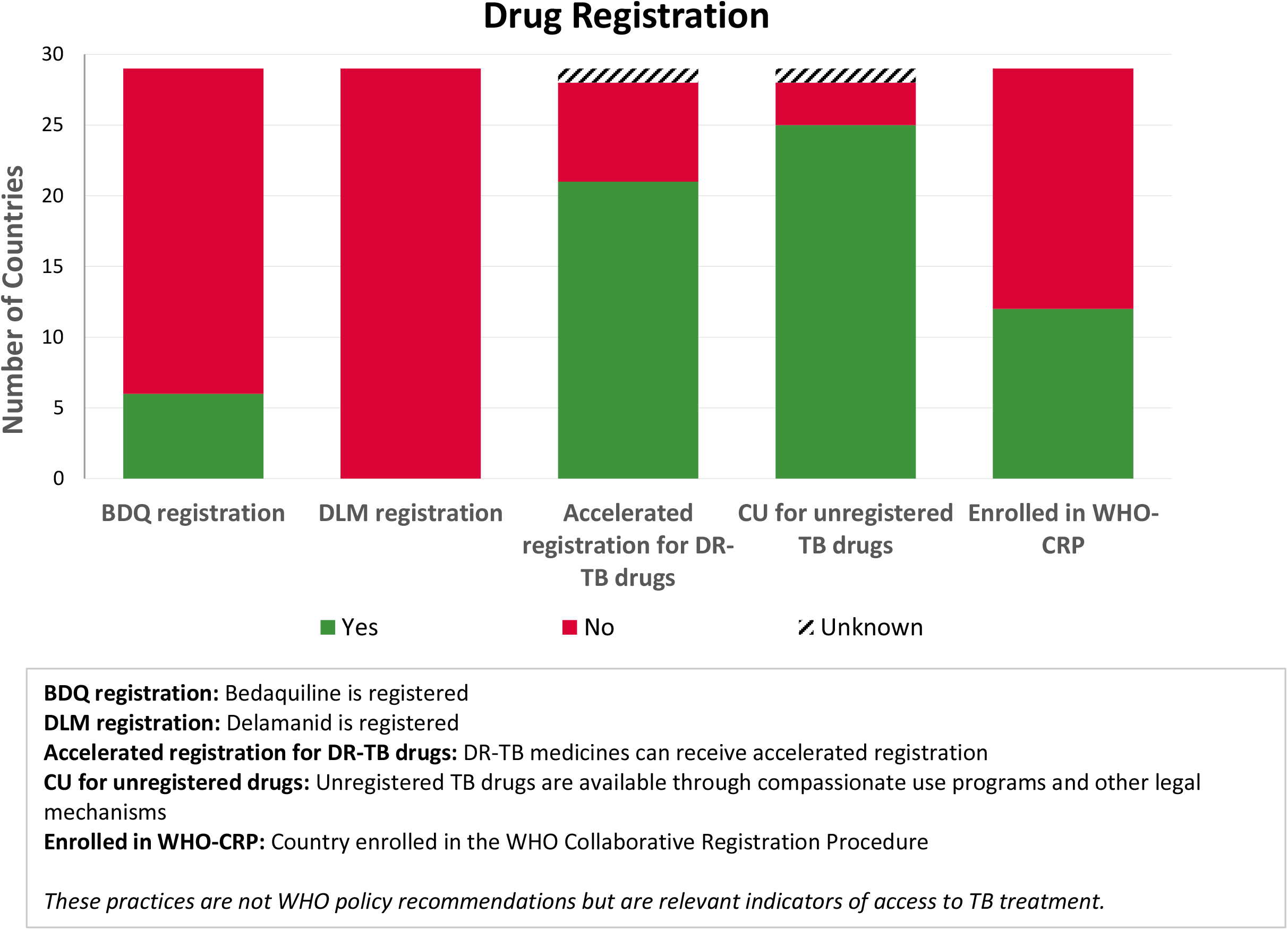
Drug registration practices adoption

## DISCUSSION

These results illustrate that many countries’ TB guidelines were not aligned with WHO recommendations at the time of the survey. While some progress had been made in the last few years, a number of countries’ TB guidelines lagged behind in key critical areas. While innovations such as the Xpert MTB/RIF test and newer drugs BDQ and DLM offer new hope for the diagnosis and treatment of TB, a lack of adoption and implementation at national level misses opportunities to reduce TB infections and deaths and hinders progress towards ending TB by 2030.

The Xpert MTB/RIF-resistance test, and the next-generation Xpert MTB/RIF Ultra test, are WHO-recommended rapid molecular tests that diagnose TB and detect rifampicin resistance in less than two hours.^30^ Until Xpert MTB/RIF entered the market in 2010, diagnosing TB relied on traditional sputum smear microscopy, which detected only half of all TB cases and could not diagnose drug resistance. Our findings showed that adoption of the Xpert MTB/RIF test has been slow: seven years after the WHO recommendation on use of Xpert MTB/RIF, just under half of the countries surveyed had yet to adopt it as an initial test for TB. Widespread use of this test has the potential to drastically reduce the TB diagnostic gap, get more people on appropriate treatment, and prevent further transmission. Others have reported significant financial and logistical challenges with the use and scale-up of Xpert MTB/RIF.^31, 32^ Countries need to address the specific challenges they face to increase access to this diagnostic tool.

Community-based, decentralized models of care enable people with TB to access care close to where they live and are associated with high treatment initiation and reduction in time to treatment;^33^ improved treatment outcomes including survival;^34^ and cost-effectiveness.^35, 36, 37, 38^ Compared with hospitalization, studies have found that hospital-based treatment does not result in better outcomes than community-based treatment for DR-TB.^39, 40^ MSF experience in South Africa showed DR-TB treatment initiation at the district level resulted in high treatment initiation and reduced time to treatment in patients.^41^ With only one in five people with DR-TB completing the diagnostic and treatment pathway in 2015, recommending the initiation of TB treatment at the community level presents an opportunity to improve this outlook.

In addition, WHO has long recognized the benefits of providing TB care integrated with HIV care,^42^ and in 2012 recommended the provision of TB and HIV services at the same location and time, i.e., a “one-stop shop” for diagnosis and treatment.^43^ The need to expand collaborative TB/HIV activities is further emphasized in the WHO End TB Strategy and should be prioritized by countries in which TB and HIV are dual public health concerns.

The results are in line with pre-existing reports of the slow adoption and implementation of the use of BDQ and DLM,^44^ demonstrating that relatively few MDR-TB patients who could benefit from these newer drugs are receiving them. In 2016, just over 4,300 people received BDQ, and only 469 people received DLM outside of clinical trials or compassionate use programs.^45^ Despite this, the majority of people with MDR-TB are left to unnecessarily endure treatment with older, more toxic and less effective treatment options. MSF is working with national programs to offer BDQ and/or DLM to people with no other treatment options, and has provided programmatic evidence showing that the combined use of these drugs provides strong signs of effectiveness for patients with few remaining options.^46^ Countries can procure both these newer drugs from Stop TB Partnership’s Global Drug Facility.^47^ Given the poor treatment success rates and high toxicity of existing regimens for MDR-TB, it is critical that countries quickly adopt and widely implement international guidelines on the use of BDQ and DLM.

Shorter treatment regimens are also urgently needed. Today, people with TB continue to face long, arduous treatment courses that can last for up to two years and require tens of thousands of pills and injections. At the time of the survey, short-course 9-month regimen was recommended by WHO. MSF showed that patients treated with a 9-month regimen had treatment outcomes similar to current standards of care.^48^ Modelling the impact of shorter treatment on MDR-TB incidence found that it has the potential to markedly reduce the incidence of MDR-TB if it was to be expanded in the absence of additional drug resistance.^49^

Although not part of a specific WHO recommendation, we looked at national drug regulation practices in this study because they have a direct effect on country-level access to TB medicines. We assessed national registrations of BDQ and DLM, as well as the availability of compassionate use programs, accelerated registration, or utilization of the WHO CRP for unregistered drugs. Not presented here, the 2017 OOS survey also examined whether or not all WHO-recommended TB medicines were included on national Essential Medicine Lists, which can help ease drug importation.^17^ Alternative access mechanisms may become increasingly important to avoid country-level shortages or stock-outs of TB drugs amid worrying global shifts in funding and support.^50^

Our survey possesses a number of caveats that must be considered regarding methodology and results analysis. In terms of country inclusion in the survey, a few high-burden countries were ruled out, due in part to poor feasibility of conducting in-depth, detailed analysis in countries where MSF or STBP had no programmatic presence. Another limitation was that the survey did not provide definitions for the level of policy implementation, be it minimal, modest, or widespread; this may have led to over-or under-reporting by countries as to the breadth of policy implementation.

During the survey period, several countries were in the process of updating their national TB guidelines. In these cases, responses were based on a mixture between completed updated guidance or pre-existing guidance where new guidance was not complete. The data presented here are accurate only for the period of the survey, and any updates in national policy adoption since this time period have not been taken into account. The time lag between recent WHO policy updates and their adoption by countries should be carefully considered when interpreting these data, as our survey was carried out through mid-2017 before the release of new or upcoming WHO guidelines in 2018. More comprehensive new TB guidelines are expected to be released by WHO near the end of 2018 or in early 2019.

## CONCLUSIONS

Looking ahead based on these findings, we hope the pace of implementation of improvements in TB care, including newer WHO recommendations that have come out since the survey, will increase more than they have in the past. The factors affecting why countries may be slow to adopt new policies and implement them for certain TB care recommendations is an area worthy of further research and analysis. While the lack of research in critical areas itself needs to be addressed, countries need to be bold and rapidly adopt available WHO guidance in these areas. The WHO itself stated in its August 2018 update on MDR-TB treatment, “While understanding that it would not be immediately possible to achieve the new standards of care in every individual MDR-TB patient, strategic planning should start immediately to enable rapid transition to the upcoming new WHO guidelines.”^27^ Not doing so will continue to unnecessarily leave thousands of people affected by TB without access to new health tools that could save their lives.

For any hope to achieve global TB health targets set by the UN High-Level Meeting on TB Political Declaration and UN Sustainable Development Goals, greater political will, funding, and resources are needed to implement the best possible TB care strategies and tools. Also, research and development into new, more effective and appropriate tests and treatments is urgently needed. If countries make bold, impactful efforts to fall in step with the latest international TB policies, we may finally put ourselves on track to see an end to the global TB crisis.

## ACKNOWLEDGMENTS

The authors would like to thank the Médecins Sans Frontières TB project teams, Stop TB Partnership members and partners, national TB programs and managers, and people affected by TB in all the countries who provided the information on which the survey was based. The authors also thank Manuela Rehr for analysis of the diagnostics indicators and results; Tracy Swan for medical writing; Hope Arcuri, Katie Glockner, Madeleine Lambert, Liam Kavanagh, Eduardo Piqueiras, and Sonya Stanczyk for help with data collection and organization, and producing the table and figures; Lucas Graf for help with editing the figures; and Oliver Yun for editorial support and manuscript coordination.

---

**S1 Text**. Supplementary methods

