## Supplementary material for "Countries are out of step with international recommendations for tuberculosis testing, treatment, and care: Findings from a 29-country survey of policy adoption and implementation": S1 Text. Supplementary methods

Supporting Information

**S1 Text. Supplementary methods**

### Countries surveyed for the Out of Step reports

The first edition of the Out of Step survey reports, published in 2014 by Médecins Sans Frontières (MSF),[1] monitored progress in the adoption of World Health Organization (WHO) international tuberculosis (TB) recommendations into national TB policies and practices in eight countries. The second edition, published in 2015 by MSF,[2] included 24 countries. The third edition, published in 2017 by MSF together with the Stop TB Partnership (STBP),[3] focused on adoption as well as implementation of policies and related practices in 29 countries.

Countries were selected for inclusion in the surveys if they had a high burden of TB, multidrug-resistant (MDR-)TB, or TB/HIV co-infection as defined by the WHO at the time,[4,5] as well as the presence of either a Stop TB Partnership member or a MSF TB project that could complete the survey. The selected countries had a wide geographical scope and economic status.

The 2017 survey included 23 of the 24 countries in the 2015 report, plus six additional countries: Armenia, Afghanistan, Bangladesh, Belarus, Brazil, Cambodia, Central African Republic (CAR), China, Democratic Republic of Congo (DRC), Ethiopia, Georgia, India, Indonesia, Kazakhstan, Kenya, Kyrgyzstan, Mozambique, Myanmar, Nigeria, Pakistan, Papua New Guinea (PNG), Philippines, Russian Federation, South Africa, Swaziland (later named Eswatini), Tajikistan, Vietnam, Ukraine, and Zimbabwe.

Twenty-six of the 29 countries in the 2017 survey had a high burden of TB, MDR-TB, and/or TB/HIV, according to WHO criteria,[5] defined as follows: For 2016-2020, each WHO high-burden country list (for TB, MDR-TB, and TB/HIV) are defined as the top 20 in terms of absolute numbers of cases plus the additional 10 countries with the most severe burden in terms of case rates per capita that do not already appear in the “top 20” and that meet a minimum threshold in terms of absolute numbers of cases (10,000 per year for TB, and 1,000 per year for MDR-TB and TB/HIV). Twelve of the 29 countries had a high burden of TB, MDR-TB, and TB/HIV; five had a high burden of TB and MDR-TB; two had a high burden of TB and TB/HIV; five had a high burden of MDR-TB; one had a high burden of TB; and one had a high burden of TB/HIV.[5] The three remaining countries (Afghanistan, Armenia, Georgia) were not included in any of the 2016-2020 WHO high burden country lists, but all were included in the lists up until 2015.[4]

While 32 countries were originally contacted for the 2017 survey, two did not respond, and one requested not to be included in the final report after completing the questionnaire.

### Development of the questionnaire

For the survey, experts from MSF and STBP developed a semi-structured questionnaire to assess the national adoption and implementation of WHO TB policies and related practices. The questionnaire included questions on TB diagnostics, models of care, treatment for drug-sensitive and drug-resistant TB, and the TB drug regulatory environment. Based on feedback from 2014 and 2015 respondents, some of the questions were rephrased in the 2015 and 2017 questionnaires, respectively, to improve clarity. The 2017 survey also included questions on TB prevention policies. The 2017 questionnaire was developed between September and November 2016.

The survey questionnaire consisted of two main parts. The first part asked whether national policies were aligned with (had adopted) current WHO policy recommendations, calling for yes or no answers. If there was no response, or an answer could not be clarified, the answer was recorded as "unknown." If the answer for policy adoption was "yes", then the second part of the questionnaire asked about implementation of these policies, asking whether and how widely they had been implemented. In this section, there were three possible responses: "yes", "yes but not widely", or "no". If the answer for policy adoption was "no", any responses about implementation were not recorded. Each question included a box asking respondents to identify the specific policy document used to support their answer. Respondents could also further comment or explain their answer in a separate "note" box. A limitation of the survey was that we did not provide definitions for the levels of implementation that the respondent could refer to (i.e., minimal, modest, or wide implementation, and associated definitions for each). Because of this limitation, we pooled implementation data, regardless of level of implementation, in the survey analysis presented here.

**Data collection**

From October to mid-November 2016, MSF and STBP conducted a desktop review of current TB and HIV policy documents and guidelines from each country (national Ministry of Health-approved policy documents and guidelines covering the relevant areas). All policy documents that could be found using online sources, as well as documents used in the 2015 survey, were compiled. In early November 2016, STBP sent an email to the national TB programs (NTPs) of all 32 countries, introducing the research, and asking them to review the documents for their country. NTPs were given approximately three weeks to confirm if these documents corresponded to the latest policy guidance being used, or to send any additional documentation.

Twenty-two of the 32 countries responded and approved the existing documents or provided additional documents. One country replied stating they were in the process of updating their guidelines and could not share their policy documents until they were completed. Countries that did not send additional documents were sent requests and reminders via email and phone calls. Nine countries did not respond after multiple attempts to contact the NTPs. Of these, two were excluded as there was no MSF in-country presence to support in completing the questionnaire without the NTP. The remaining seven countries had an MSF project presence in the country, allowing the MSF teams there to complete the questionnaire.

This process of collecting policy documents continued until the end of December 2016. Where source documents were only available in the local language, these were translated by a professional translation service. The full list of documents obtained is available at: www.stepupfortb.org.

*Completing the questionnaires: countries with MSF presence*

MSF followed up with the 19 countries where it operates TB projects. One country asked not to be included in the survey after completing the questionnaire but before the report went to print. The remaining 18 MSF countries were: Armenia, Belarus, Brazil, CAR, DRC, Georgia, India, Kenya, Kyrgyzstan, Mozambique, Myanmar, PNG, Russian Federation, South Africa, Swaziland (later named Eswatini), Tajikistan, Ukraine, and Zimbabwe.

MSF project teams in each country were contacted to complete the questionnaire. The questionnaires were shared in December 2016 for completion by 20 January 2017. All the projects completed and returned the questionnaires. Once completed, the questionnaires were sent to a consultant fact checker and to a representative from STBP who reviewed the responses to ensure they reflected the policies in the documents that had been collected, or that had been shared by the projects. If one or more of the documents mentioned in the responses was missing, and we could not identify the source of the information, we sought clarification with the MSF project by email or phone calls.

*Completing the questionnaires: countries with STBP presence*

STBP followed up in 11 countries: Afghanistan, Bangladesh, Cambodia, China, Ethiopia, Indonesia, Kazakhstan, Nigeria, Pakistan, Philippines, and Vietnam. Using the policy documents that had been collected, STBP pre-filled the questionnaire. One country was in the process of updating their guidelines and had not shared their policy documents, so it was not possible to pre-fill their questionnaire; they completed the questionnaire themselves based on a combination of the completed updated policies and pre-existing guidelines.

In some cases, the responses were checked by a second colleague at STBP, as well as the consultant fact checker, and then sent to the 11 NTPs for verification. In other cases, after being pre-filled, the questionnaire was sent directly to NTPs, and then checked by a second STBP colleague and the consultant fact checker after verification by the NTPs. If there were any discrepancies, or we could not locate a document that the NTP referred to, we communicated with the NTPs for clarification. The questionnaire was translated from English to Russian for one NTP.

Ten NTPs validated the pre-filled questionnaires. One country did not validate the pre-filled questionnaire, but they were included in the report with the lack of NTP validation clearly noted. The validation process began in February 2017 and was completed in mid-May 2017.

During the survey period, some countries were in the process of updating their guidelines. In these cases, responses were based on a combination of updated guidance and pre-existing guidance where new guidance was not complete. The time lag between recent WHO policy updates and their adoption by countries should be taken into account when analyzing the data. Although originally included in the questionnaire, questions about policies and implementation of TB-LAMP (loop-mediated isothermal amplification) were not included in the survey, since the guidance was very new when the data collection began.

**Data validation**

Once the questionnaires were completed, the relevant sections in each area – diagnostics, models of care, treatment, prevention, and drug regulatory environment – were reviewed by MSF experts. Their feedback was sent to a consultant fact checker, who then led a process of working with the experts to re-contact MSF projects to clarify responses for MSF-project countries where necessary, while STBP re-contacted the NTPs of the countries where STBP was present.

**Data analysis**

Key findings were provided for each area, reported as both percentages and numbers. Unless noted, the denominator was 29, for all countries included in the survey. If a country did not answer a question, both the numerator and denominator were adjusted. Comparisons and progress made from the 2015 survey to the 2017 survey were reported where possible for the 23 countries included in both surveys, reported as both percentages and numbers. Only data that could be verified with no discrepancies were taken into account for this comparison.

The full results of the survey can be found in the OOS 2017 report.[3]
